# Identifying pyrogenic contaminants using transcriptomic profiling of monocyte activation test with machine learning

**DOI:** 10.1101/2025.08.13.670109

**Authors:** Tess AV Afanasyeva, Bruno FM de Albuquerque, Paulien Doodeman, Miranda C Dieker-Meijer, Marijke Molenaar-de Backer, Teunis JP van Dam, Anja ten Brinke

**Author notes:** These authors contributed equally to this work (Teunis JP van Dam, Anja ten Brinke).

## Abstract

The monocyte activation test is an *in vitro* pyrogenicity assessment method that can utilise human peripheral blood mononuclear cells to detect pyrogens in injectable drugs, providing a binary outcome that indicates the presence or absence of a pyrogen. The added ability to distinguish between different types of pyrogens would greatly expand the applicability of the test, for example, by allowing to pinpoint the source of a contaminating pyrogen in pharmaceutical products. Pyrogens activate a unique set of pattern recognition receptors (PRRs), which contribute to inflammation, yielding distinct transcriptomic activation signatures. In this paper, we capture the unique expression signatures of activated monocytes through bulk RNA sequencing and introduce a data preprocessing pipeline that allows the training of a machine-learning model to classify pyrogenic contaminants. Using a dataset of 108 samples stimulated with five classes of PRR agonists, we could differentiate between these classes with more than 97% F1 on test data. We further demonstrate the model’s capacity to generalise on the previously unseen data using different ligands for the same PRRs as well as heat-killed *Escherichia coli* and *Staphylococcus aureus*.

## INTRODUCTION

Pharmaceutical products intended for parenteral administration must comply with stringent safety standards to ensure the absence of pyrogenic contamination, which can induce systemic inflammation, fever, and, in severe cases, septic shock and even death. Contaminating pyrogens encompass a range of substances, including endotoxins derived from Gram-negative bacteria and non-endotoxin pyrogens from Gram-positive bacteria, viruses, and fungi. In response to these pyrogens, human monocytes produce pro-inflammatory cytokines, such as cytokine IL-6, a phenomenon leveraged in the in vitro monocyte activation test (MAT).

The MAT is an animal material-free method that relies on the detection of pro-inflammatory IL-6 using enzyme-linked Immunosorbent Assay (ELISA). It consists of human peripheral blood mononuclear cells (PBMCs) isolated from the buffy coat of four human volunteers, pooled, and cryopreserved. The MAT has been demonstrated to have high sensitivity and specificity in detecting both endotoxin and non-endotoxin pyrogens (Daniels, 2022; Hoffmann et al., 2005; Solati et al., 2015). The current state-of-the-art MAT provides either a pass/fail result, indicating whether PBMCs produce IL-6 or not, or, in cases when the product dilutions align with the standard endotoxin range, a semi-quantitative pyrogen concentration readout in biological activity units, called endotoxin equivalents per millilitre (EE/mL) (chapter 2.6.30, European Pharmacopoeia, 2025). Since all pyrogens induce IL-6 production, the current MAT is not able to distinguish between types of pyrogens.

Insights into the type of contaminating pyrogens in parenteral pharmaceutical products can prevent future contaminations and lead to improvements in manufacturing procedures and product safety. Currently, the identification of product-contaminating pyrogens is conducted through targeted screening with specialized reporter cell lines expressing a single type of Pattern Recognition Receptors (PRRs) (Hacine-Gherbi et al., 2017; Huang et al., 2009). This approach depends on the availability of commercial reporter cell lines and is thus limited in PRR coverage. Furthermore, some pyrogens activate multiple PRRs simultaneously, leading to incomplete characterisation with a single PRR receptor cell line. Here, we tackle this challenge by developing a computational model to analyse the transcriptomic profiles of pyrogen-stimulated PBMCs, the unique expression signatures of which can be captured through bulk RNA sequencing (Salyer & David, 2018). The significant overlap in gene expression between the activated pathways makes it challenging to distinguish the activating pyrogens by standalone statistical methods such as principal component analysis (PCA). Therefore, we employ a machine-learning (ML) classification approach, where we generated 108 MAT samples exposed to either Toll-like receptor (TLR1/2, TL4, TLR5, TLR7/8), or NOD2 activator ligands and train ML models with an F1 score exceeding 97% on test data. We further demonstrate the model’s generalisation capacity using RNA sequencing data of ligands not trained on, as well as heat-killed *Escherichia coli (E.coli)* and *Staphylococcus aureus (S. aureus)*.

## METHODS AND MATERIALS

### Monocyte Activation Test Assay

The MAT was performed using the concentrations of ligands as shown in Table 1 and 1 lot of MAT Cell Set HS (Essange Reagents, The Netherlands) following the manufacturer’s instructions with modifications (Solati et al., 2015). In short, a vial containing ∼5×10^6^ (± 1×10^6^) cryopreserved PBMCs of a pool of 4 donors (1 ml) was thawed in a water bath at 37°C, washed with in Iscove’s Modified Dulbecco’s Media supplemented with 5% human serum and taken up in the same medium to obtain a final cell concentration of 200,000 cells per well (Lonza, Belgium or ThermoFisher, US) (Essange Reagents, The Netherlands) (Molenaar-De Backer et al., 2021). Each ligand was titrated to determine the concentration that would produce an equivalent amount of IL-6 protein to 0.16 Endotoxin Units/ml of LPS under the same conditions. IL-6 protein was measured with the IL-6 ELISA Kit (Essange Reagents, The Netherlands) according to the manufacturer’s instructions. MAT cells (200,000 cells per well) were incubated at 37°C at 5% CO_2_ for 3.5 hours. MAT was performed with 0.16 EE/ml of each ligand, cells with culture medium only was used as negative control. After 3.5 hours incubation total RNA was isolated using RNeasy mini kit on QIAcube Connect, as per the manufacturer’s instructions, followed by DNAse I treatment (Qiagen, Netherlands). As a quality control step, IL-6 protein expression levels were measured in duplicate in the supernatants of these samples using the IL-6 ELISA Kit (Essange Reagents, The Netherlands).

**Table 1.** Overview of pyrogenic ligands, their concentrations used, PRR targets, and sample size. LPS was Biological reference preparation from EDQM, the rest of the ligands were obtained from Invivogen, San Diego, CA, USA.

| Ligand | Concentration<br>used | Activates<br>TLR | Sample<br>size | Train<br>set? |
| --- | --- | --- | --- | --- |
| IMDM only (Negative Control) | none | none | 21 | T |
| Lipopolysaccharides (LPS, *E. coli*<br>0113:H10) | 0.16 EU/mL | TLR4 | 22 | T |
| Flagellin (Fla-Pa, *P. aeruginosa*) | 20 ng/mL | TLR5 | 13 | T |
| Peptidoglycan (PGN) | 30 µg/mL | TLR2 | 31 | T |
| Pam3CSK4 (Pam3) | 2.5 ng/mL | TLR1/2 | 16 | T |
| Resiquimod (R848) | 2 µg/mL | TLR7/8 | 5 | T |
| Pam2CSK4 (Pam2) | 10 ng/mL | TLR2/6 | 6 | F |
| Lipoteichoic Acid (LTA) | 60 ng/mL | TLR2 | 3 | F |
| Monophosphoryl Lipid A (MPLA) | 2 µg/mL | TLR4 | 4 | F |
| Heat-Killed *S. aureus* | 5×10<sup>6</sup> cells/mL | multiple | 6 | F |
| Heat-Killed *E. coli* | 5×10<sup>4</sup> cells/mL | multiple | 6 | F |
| Total |  |  | 133 |  |
| Total Training Set |  |  | 108 |  |

### HEK-blue TLR testing

Pyrogens (at a concentration equivalent to 0.16 EU/ml of LPS and the same as in the rest of this paper) were tested using HEK-Blue™ hTLR2 cells, HEK-Blue™ hTLR4 cells, and HEK-Blue™ TLR5 cells (Invivogen). In short, 50.000 cells were plated in 150 µL HEK-Blue™ Detection medium (InvivoGen) in a flat-bottom 96-well culture plate to which 50 µl sample or standard curve was added. For HEK-Blue™ hTLR4 cells, 10 µL of FBS and 40 µL of sample or standard were added. Plates were incubated at 37°C, 5% CO2. After 24 hours, the plates were measured at 630 nm with a spectrophotometer for SEAP (Secreted Embryonic Alkaline Phosphatase) detection.

### Transcriptomic Analysis

The RNA was sequenced using poly(A) enriched standard library preparation using NEBNext Ultra II Directional RNA Library Prep Kit (New England Biolabs, USA). Paired-end 150-bp sequencing was performed on Illumina NovaSeq 6000 instrument (GenomeScan B.V.), with sequence coverage of 20 million reads. Unique Molecular Identifiers (UMIs) were used to address PCR duplication (Kivioja et al., 2012). The samples that did not pass quality control were excluded from further analysis. RNA-Seq data processing was conducted using a custom Snakemake (v8.23.1) pipeline. First, raw reads were trimmed and filtered using fastp (v0.23.4). The poly-G tails, low-quality bases, and adaptors were removed, low-complexity reads were filtered out and reads longer than 36 base pairs were retained. Next, reads were aligned to the GRCh38_GCA_000001405.15 reference genome using STAR (v2.7.11a) and the resulting BAM files were then merged and reindexed using samtools (v1.19.2) to handle duplicate runs. Following this, deduplication of the aligned BAM files was performed using UMI-tools (v1.1.4), using the directional method, discarding chimeric pairs and unpaired reads. Finally, gene expression counts were generated using featureCounts (from the subread package, v2.0.6). Only fragments where both ends were mapped successfully were counted, and chimeric fragments were excluded. The counts and raw data are submitted to Gene Expression Omnibus (GEO accession number GSE313994).

### Differential Gene Expression

Gene expression data was corrected, filtered, and normalised using the PyDESeq2 (v0.4.11) (Love et al., 2014; Muzellec et al., 2023) in Python 3.10.14. Genes were filtered out by having a total sum lower than 10 counts. Differentially expressed genes across all samples were compared with the IMDM-only control samples with the FDR cut-off of 0.05 and |log 2-FC|> 2. Go terms were retrieved using goatools (Klopfenstein et al., 2018). PCA was performed using scanpy (v1.10.3) (Wolf et al., 2018). The counts and raw data have been submitted to the Gene Expression Omnibus repository (accession number GSE313994).

### Machine Learning

Data preparation and classification model testing were performed using scikit-learn (v1.5.2) (Pedregosa et al., 2011). On the total of 108 training samples, a 5-fold stratified cross-validation approach was applied, with 20% of the data held aside as a test set, while the remaining 80% was used for feature selection and model training. During preprocessing, duplicated gene counts across all samples were removed, and normalisation to library size was conducted. Genes were ranked based on their dependency with the pyrogen class variable using chi-squared test statistics, and the top 1000 genes were selected. From this set, 250 genes were further chosen using the Random Forest estimator, with importance weights above a 0.01 threshold. A total of 250 trees with a maximum depth of 5 were used. We used SMOTE from imbalanced-learn package (v0.13.0) to oversample non-majority of classes and to generate synthetic samples to bring all the classes to 20. We further standardised the transcript counts. Multiple classification models—Linear Support Vector, Linear Support Vector with stochastic gradient descent (SGD), Random Forest (all from scikit-learn (v1.5.2)), and XGBoost (v2.1.1)—were compared using accuracy, precision, recall, F1-score. Accuracy is measured by calculating the ratio of correct predictions (true positives and true negatives) to the total predictions made. Precision is determined by calculating the fraction of true positives versus all positives, while recall is determined by calculating the fraction of true positives out of all positive predictions. The F1-score balances precision and recall by calculating the harmonic mean. The code has been made available on github.com/sqn-bioinformatics/MATseq.

## RESULTS

### Pyrogen Selection and Titration

The MAT detects pyrogenic contamination in pharmaceutical products using cryopreserved pooled human PBMCs by determining IL-6 protein production in the culture supernatant. Using the same PBMCs, here we expand the MAT capacity for the characterisation of pyrogenic contaminants using RNA expression profiles (Figure 1). The pooled PBMCs of four donors were stimulated with purified or synthetic TLR ligands, and their responses were analysed. To obtain enough RNA the number of PBMCs per well was increased from 50,000 cell per well for MAT to 200,000 cells per well for sequencing. Firstly, we titrated the pyrogens to obtain comparable concentrations in the range of 0.16 EE/ml, which is equivalent to the range of concentrations of contaminations observed in real-world samples tested at Sanquin’s MAT services laboratory. As a positive control, we used TLR4 activator LPS also called endotoxin (Findlay et al., 2015; Poltorak et al., 1998). Additional ligands were selected to represent a diverse group of TLR activators, such as TLR2/TLR1 heterodimers activator synthetic triacylated lipopeptide Pam3 (Pam3CSK), NOD2 activator bacterial peptidoglycan PGN, TLR5 activator flagellin, and R848, a synthetic imidazoquinoline compound, which stimulates both TLR7 and TLR8 to elicit a virus-like response (Aliprantis et al., 1999; Chuang & Ulevitch, 2000; Girardin et al., 2003; Hayashi et al., 2001). A dataset of total size of 133 samples was used for the analysis, of which 108 samples belonged to the training set (Table 1).

**Figure 1.**
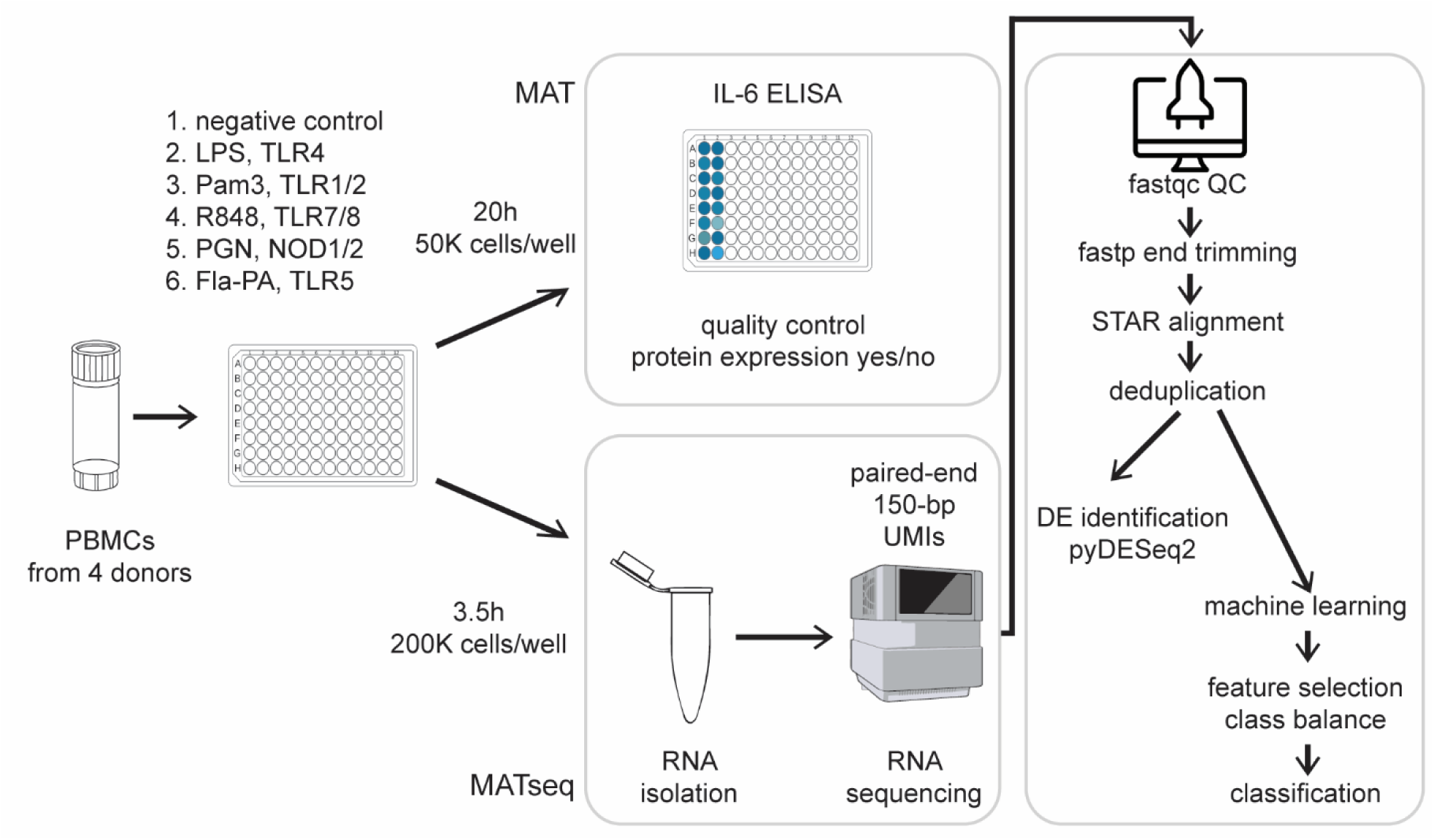
Graphical summary of the assay set-up in this study.

### Differential Gene Expression Analysis

Differential gene analysis showed an upregulation of the immunology-related genes in PBMCs stimulated with each class of PRR activators. We observed upregulation of expected inflammatory mediators such as IL6, as well as multiple chemokines, for example, CCL3 and CCL4 (Figure 2, Supplementary Table 1). The following GO terms were consistently enriched across all ligands (Fla-PA, LPS, PGN, Pam3, R848): inflammatory response (GO:0006954), immune response (GO:0006955), cellular response to LPS (GO:0071222), and cytokine-mediated signalling pathway (GO:0019221). R848 stimulation induced an antiviral and interferon-mediated profile such as defence response to the virus (GO:0051607), antiviral innate immune response (GO:0140374), and type I interferon-mediated signalling pathway (GO:0060337)(Supplementary Table 2).

**Figure 2.**
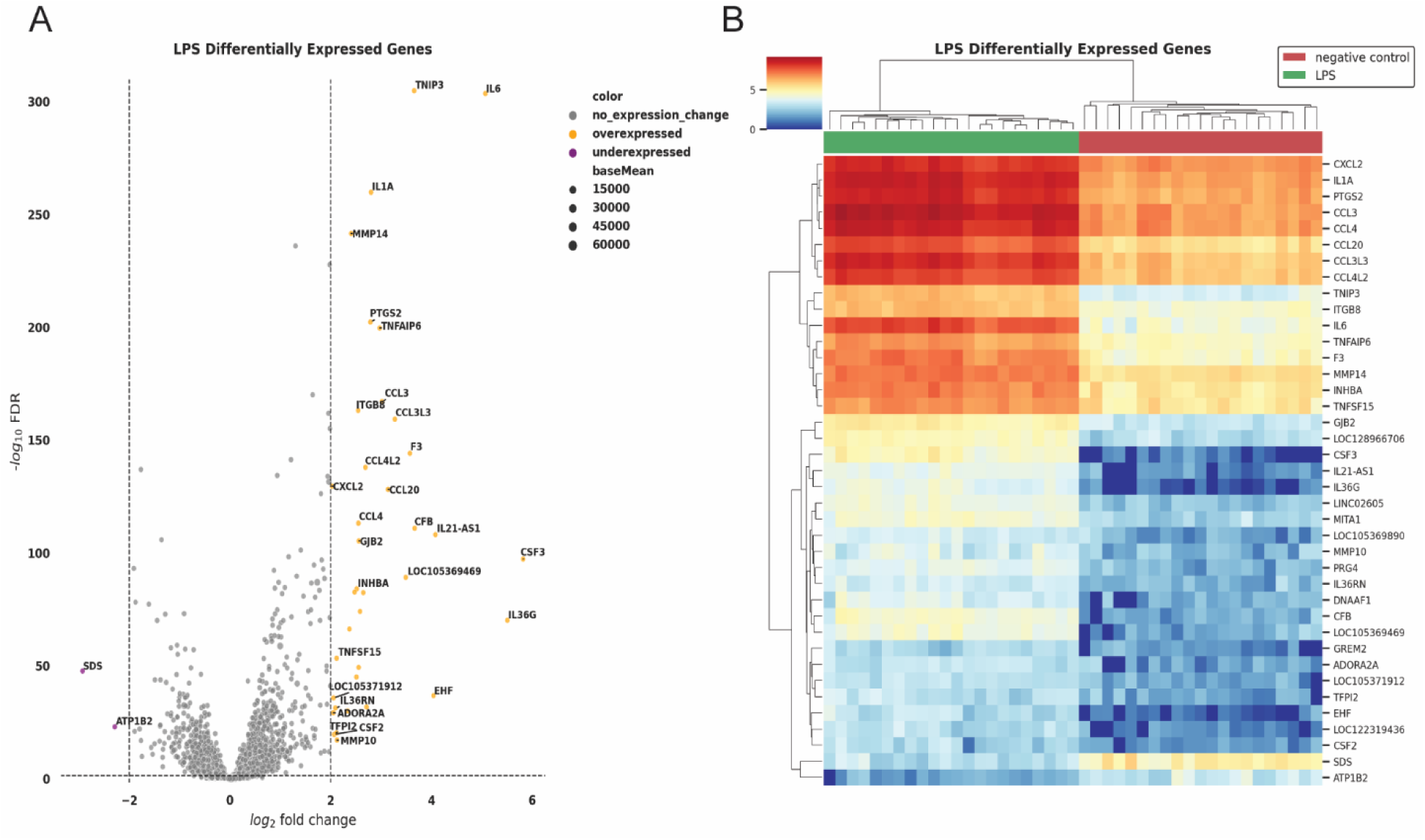
A representative example of differential gene expression analysis for ligand-stimulated PBMCs. A) Up-and down-regulated genes for LPS-stimulated cells (FDR = 0.05, log2FoldChange =2, B) Differential gene expression for LPS-stimulated PBMCs.

### Data Preprocessing

The PCA of the transcripts of the PBMCs stimulated with different classes of ligands showed no distinguishable separation based on the ligand class. Therefore, we applied feature selection pipeline to select the most relevant genes to improve class separation and classification (Figure 3A,B). Firstly, genes with gene counts that were the same across all samples were dropped. This step removes genes with zero counts across all samples and is performed before the normalisation step to decrease the computational and time resources required for the analysis. Further, gene counts were normalised to the library size of the respective sample. Next, we selected 1000 gene transcripts with the largest differences in distribution frequencies between ligand classes using chi-squared test. We fitted several randomised decision trees to select 250 genes that performed best for each run. When comparing PCA on either pre and post-feature selection, we observed an increased degree of separation of the different ligand classes (Figure 3B). Interestingly, Fla-PA clustered with Pam3 and LPS classes, depending on experimental date and batch number.

**Figure 3.**
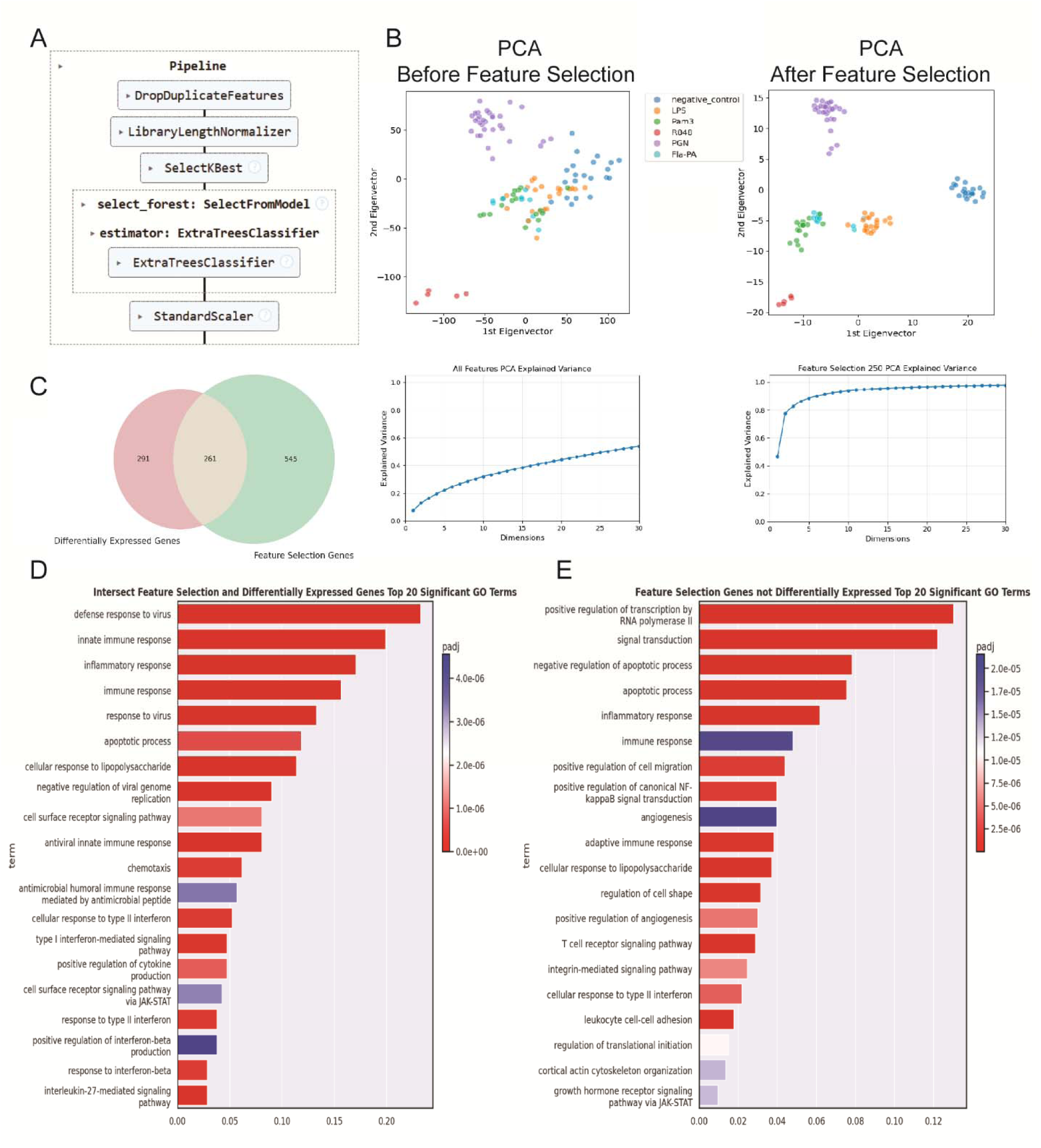
Feature selection pipeline improves the separation between classes of transcriptomic activation profiles in PBMCs. A) Steps of the feature selection pipeline. Genes with no expression change across all samples were removed with DropDuplicateFeatures, and the remainder was normalised to library size using LibraryLengthNormalizer. SelectKBest was used to select the top 1000 features using the chi-squared test and Extra Trees Classifier to choose the top 250 of the most informative genes. B) PCA before and after the feature selection showed improved clustering of the classes based on the class of stimulating ligand. C) Overlap of genes post feature selection (n=1000 runs at 80% of all samples) with differentially expressed genes consisting of 261 genes. D) The genes post feature selection belong to biological processes GO terms related to topics inflammation, innate immunity, and inflammatory response. E) The genes post feature selection that are not differentially expressed belong to biological processes GO terms related to topics RNA polymerase II upregulation, signal transduction, apoptosis, but also inflammatory response, cellular response to lipopolysaccharide and immune response.

To obtain insights in which genes are being selected as relevant, we run the feature selection pipeline 1000 times on randomly selected 80% of the training data. We observed genes *IL6, C5AR, TGFBI, PTGS2, MPP1, CXCL8, CSF1R, TNFSF8, CXCL2, PELI1, ENG*, and *PECAM1* to be present in all runs, all of which, except for *MPP1, TNFSF8*, and *PELI1*, were also found to be differentially expressed in at least one of the classes (Supplementary Table 3). Overlap of genes post feature selection of the training set (n=1000, with different random seeds) with differentially expressed genes consisting of 261 genes (Figure 3C). These intersect genes belonged to biological processes GO terms related to topics inflammation, innate immunity, and inflammatory response, however, the abovementioned GO terms were also present in the post feature selection genes that were not differentially expressed (Figure3D,E, Supplementary Table 4).

### Model Training

We compared classification performance of ML models with all gene counts/features, with 250 randomly selected genes, with 250 most informative genes selected by the feature selection pipeline, and with 250 genes added together with the differentially expressed genes. We compared Linear Support Vector Classifier (SVC), Linear Support Vector Machine with stochastic gradient descent (SGD) learning (Vapnik, 2000), Random Forest (Breiman, 2001), and XGBoost (Chen & Guestrin, 2016). The Linear SVC achieved perfect accuracy (1.0) and F1 scores after feature selection (Table 2, all numbers are shown on the test set). Random Forest also performed well after feature selection, reaching 1.0 accuracy and F1 scores. Performance of SGD improved after feature selection from under 0.89 of F1 score to 1.0. Surprisingly, XGBoost performed worse than other models both at all genes and after feature selection. As expected, all models performed worse when trained on randomly selected genes than on post-feature selection genes. Interestingly, when we added genes that were found to be differentially expressed between each pair of the ligand classes to the set of genes that were selected by the feature selection pipeline, we observed lowered performance for all models except for LinearSVC. We continued with a dataset of genes after feature selection pipeline in combination with either LinearSVC or with Random Forest. Overall, we increased the classification performance of simple ML models on the dataset with a small number of samples by filtering for the most informative ones.

**Table 2.**
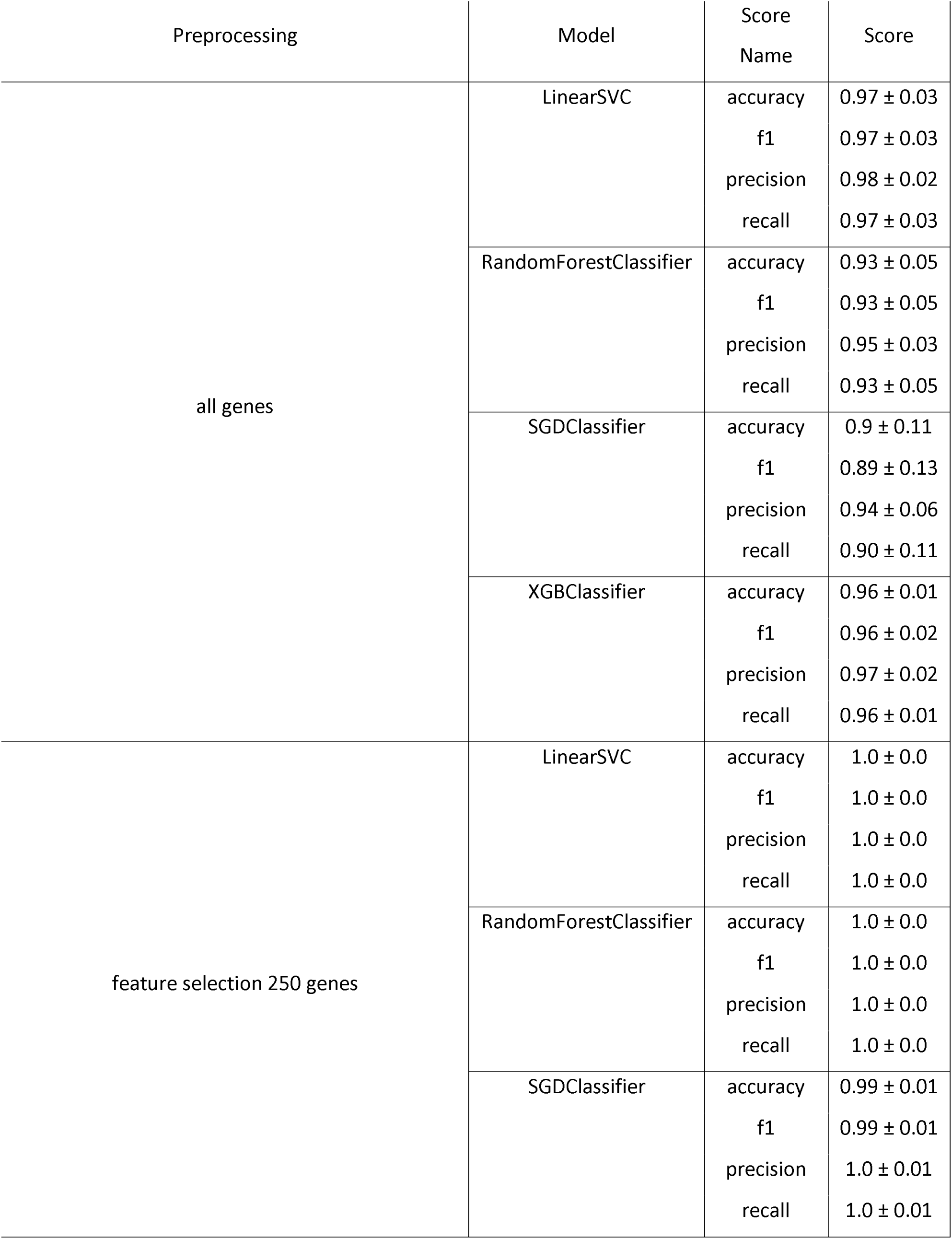
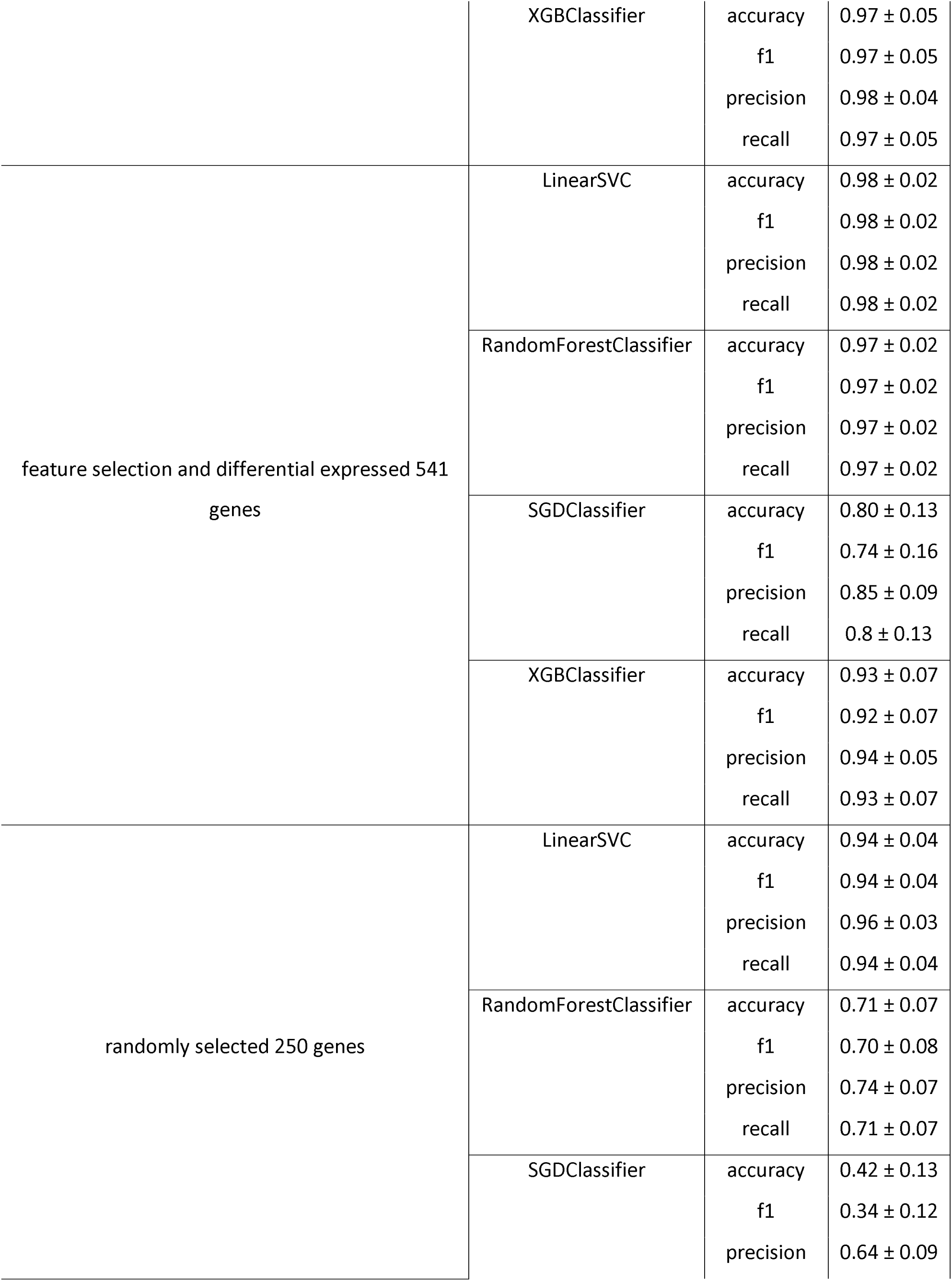
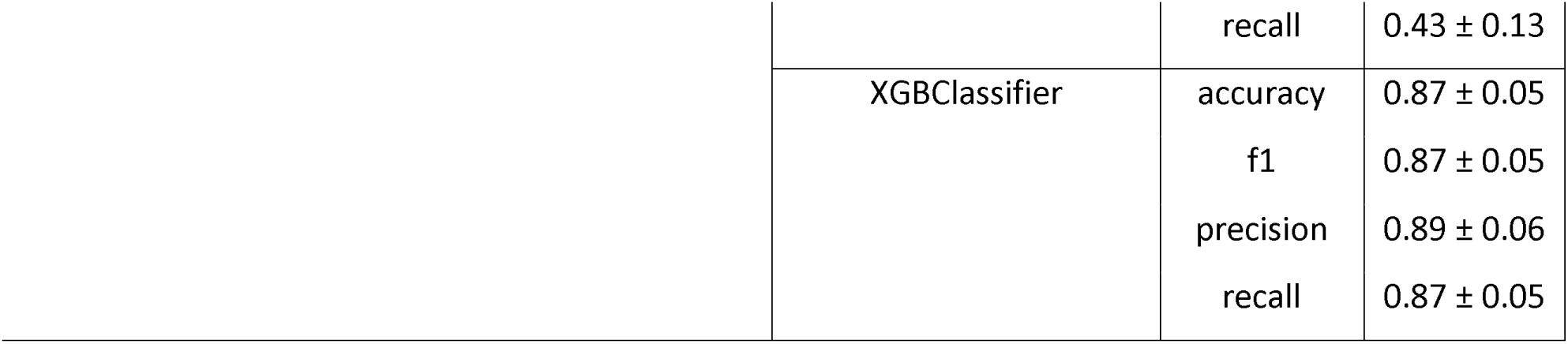
Comparison of ML models performance on test set, with all features, feature selection, both post-feature selection and differentially expressed genes, and randomly selected 250 genes where mean±standard deviation of 5 measurements are shown for accuracy, F1 score, precision and recall for Linear SVC, Random Forest, SGD, and XGBoost models on test set.

### Prediction on Ligand not in Training Set and Complex Bacterial Samples

To confirm the capacity of the tested models to generalise to previously unseen TLR activators, we generated samples with different pyrogens, distinct from those used in the training dataset but activating the same TLRs. We selected TLR2 agonist lipoteichoic acid (LTA), a major constituent of the cell wall of Gram-positive bacteria (Schwandner et al., 1999) and Pam2, a synthetic TLR2 agonist derived from bacterial lipoproteins (Parra-Izquierdo et al., 2021). In the PCA, after feature selection, these samples clustered together with Pam3, TLR 1/2 agonist, and a subset of Fla-PA, a TLR5 agonist (Figure 4A). Further, we tested MPLA, a TLR4 agonist, the immunostimulatory component of LPS from Gram-negative bacteria (Fensterheim et al., 2018). As expected, MPLA clustered with LPS on the PCA. Using the feature selected dataset and probability prediction function of either LinearSVC or Random Forest, we checked to which class their models were more likely to attribute these previously unseen ligands. Both models agreed on the most likely predicted class, which in turn, agreed with the observed PCA clustering. While MPLA is correctly predicted, LTA and Pam2 ligands were partly wrongly predicted as Fla-PA (Figure 4B,C).

**Figure 4.**
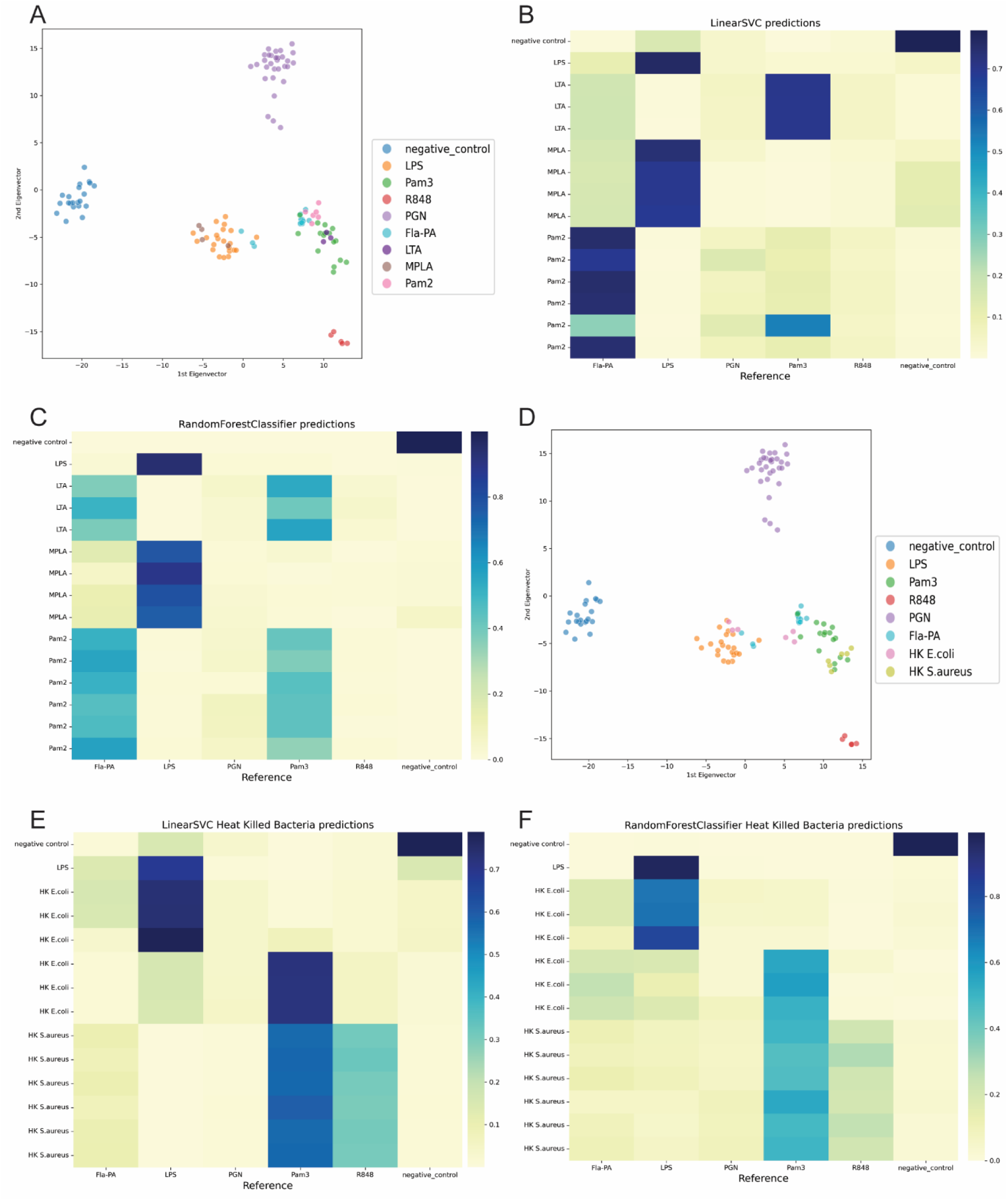
Predictions of ML models on previously unseen TLRs and heat-killed bacterial samples. A) PCA after feature selection of classes of previously unseen ligands, B,C) LinearSVC and RandomForest Classifier-based classification of previously unseen ligands, D) PCA after feature selection of heat-killed *E.coli* and *S. aureus*, E,F) LinearSVC and RandomForest Classifier-based classification of heat-killed *E.coli* and *S. aureus* post feature selection.

Furthermore, we tested heat-inactivated bacterial samples to mimic a complex bacterial contaminant. *S. aureus* and *E.coli* are expected to activate TLR2 and TLR2/TLR4, respectively. In PCA, after feature selection, *E.coli* clustered together with either LPS, a TLR4 agonist or Pam3, a TLR2 agonist, depending on to which experimental replicate samples belonged to. This is also reflected in predictions of the models, which agreed on predicting the *E.coli* samples as TLR4 or TLR2 activator depending on the experimental replicates, and *S. aureus* as Pam3 and to a lesser extent R848 (Figure 4E,F).

## DISCUSSION

In this study, we developed a ML model to classify contaminants in biological products. Our goal was to distinguish between specific PRR agonists using transcriptomic profiles of activated PBMCs. We adjusted the standard MAT to accommodate RNA sequencing requirements, titrated the ligand to standardize the endotoxin equivalents, sequenced RNA of 133 samples, developed a gene selection pipeline to improve ML model performance and sample clustering, and subsequently trained ML models to classify the PRR activators. We achieved a largest possible F1 score with 3 separate ML models on the test data. To our knowledge, this is the first study to classify contaminating pyrogens in MAT context using transcriptomic profiling and ML. Our results indicate that transcriptomic profiling can effectively distinguish between different PRR activators, suggesting its potential as a reliable method for detecting and characterizing pyrogenic contaminants.

It has been previously demonstrated that transcriptomic signatures induced by various TLR agonists can effectively discriminate between extracellular and intracellular innate immune stimuli. Specifically, Salyer and David et al. observed that extracellular TLRs (TLR2, TLR4, and TLR5) predominantly induced CXC chemokines such as *CXCL5, CXCL6*, and *CXCL8*, whereas intracellular receptors like TLR7 and TLR8 primarily upregulated *CXCL11* and *CXCL12* (Salyer & David, 2018). Similarly, we observed when selected based on their information content and usefulness for ML prediction, a subset of just 250 genes was sufficient to differentiate between classes of PRR agonists.

After feature selection, we observed preferential clustering of Fla-PA with either Pam3 or LPS in the PCA, which could influence the misprediction of the Pam3 or LPS class as Fla-PA by the ML model. Particularly, the TLR2 activators LTA and Pam2 were predicted as Fla-PA. This could be due to similarities in the induced signalling pathways, to the presence of exogenous flagellin in the samples, or due to potential contamination of the Fla-PA preparation, as previously reported (Mueller et al., 2012). Indeed, when the TLR5 activator Fla-PA was tested using the HEK-Blue™ reporter cell line system, we observed a signal for TLR2 and TLR4 (Supplementary Tables 5 and 6 and Supplementary Figure 2). Irrespective of the underlying reason, the split in the Fla-PA samples, either clustering more to Pam3 or LPS, drove the misclassification of LTA and Pam2. When we repeated the analysis excluding Fla-Pa from training classes, and observed expected classification of LTA and Pam2 samples (Supplementary Figure 3). Future work is needed to obtain definitive Fla-PA RNA transcripts.

Taken together, this study highlights the utility of ML-driven transcriptomic profiling of PBMCs for identifying pyrogenic contaminants in pharmaceutical products. However, further validation is needed with larger, more diverse datasets to ensure robustness across a broader range of pyrogenic ligands, as well as additional heat-inactivated bacterial species and elements of viral capsid to capture an expansive range of the transcriptomic responses of challenged PBMCs in MAT.

## ACKNOWLEDGEMENTS

This work was funded by Sanquin Diagnostics B.V., and Sanquin Reagents B.V, The Netherlands under PPODR20-40 and 21-03.

## CONFLICT OF INTERESTS

Sanquin Diagnostics B.V. performs MAT testing for customers. Sanquin Reagents B.V. produces and distributes MAT cell kits.

## Notes

https://www.ncbi.nlm.nih.gov/geo/query/acc.cgi?acc=GSE313994

### Competing Interest Statement

The authors have declared no competing interest.

### Summary of Updates

We updated Figure 3 and added Supplementary Figure 2.

## REFERENCES

Aliprantis, A. O., Yang, R.-B., Mark, M. R., Suggett, S., Devaux, B., Radolf, J. D., Klimpel, G. R., Godowski, P., & Zychlinsky, A. (1999). Cell Activation and Apoptosis by Bacterial Lipoproteins Through Toll-like Receptor-2. Science, 285(5428), 736–739. 10.1126/science.285.5428.736

Breiman, L. (2001). Random Forests. Machine Learning, 45(1), 5–32. 10.1023/A:1010933404324

Chen, T., & Guestrin, C. (2016). XGBoost. Proceedings of the 22nd ACM SIGKDD International Conference on Knowledge Discovery and Data Mining, 785–794. 10.1145/2939672.2939785

Chuang, T., & Ulevitch, R. (2000). Cloning and characterization of a sub-family of human toll-like receptors: hTLR7, hTLR8 and hTLR9. European Cytokine Network, 11(3), 372–378.

Daniels, R. (2022). Validation of the monocyte activation test with three therapeutic monoclonal antibodies. ALTEX. 10.14573/altex.2111301

European Pharmacopoeia. (2025). EDQM – European Directorate for the Quality of Medicines and HealthCare. In EDQM – European Directorate for the Quality of Medicines and HealthCare onocyte Activation Test Chapter: Vol. 2.6.30 (11.7th edition).

Fensterheim, B. A., Young, J. D., Luan, L., Kleinbard, R. R., Stothers, C. L., Patil, N. K., McAtee-Pereira, A. G., Guo, Y., Trenary, I., Hernandez, A., Fults, J. B., Williams, D. L., Sherwood, E. R., & Bohannon, J. K. (2018). The TLR4 Agonist Monophosphoryl Lipid A Drives Broad Resistance to Infection via Dynamic Reprogramming of Macrophage Metabolism. The Journal of Immunology, 200(11), 3777–3789. 10.4049/jimmunol.1800085

Findlay, L., Desai, T., Heath, A., Poole, S., Crivellone, M., Hauck, W., Ambrose, M., Morris, T., Daas, A., Rautmann, G., Buchheit, K. H., Spieser, J. M., & Terao, E. (2015). Collaborative study for the establishment of the WHO 3(rd) International Standard for Endotoxin, the Ph. Eur. endotoxin biological reference preparation batch 5 and the USP Reference Standard for Endotoxin Lot H0K354. Pharmeuropa Bio & Scientific Notes, 2015, 73–98.

Girardin, S. E., Travassos, L. H., Hervé, M., Blanot, D., Boneca, I. G., Philpott, D. J., Sansonetti, P. J., & Mengin-Lecreulx, D. (2003). Peptidoglycan Molecular Requirements Allowing Detection by Nod1 and Nod2. Journal of Biological Chemistry, 278(43), 41702–41708. 10.1074/jbc.M307198200

Hacine-Gherbi, H., Denys, A., Carpentier, M., Heysen, A., Duflot, P., Lanos, P., & Allain, F. (2017). Use of Toll-like receptor assays for the detection of bacterial contaminations in icodextrin batches released for peritoneal dialysis. Toxicology Reports, 4, 566–573. 10.1016/j.toxrep.2017.10.004

Hayashi, F., Smith, K. D., Ozinsky, A., Hawn, T. R., Yi, E. C., Goodlett, D. R., Eng, J. K., Akira, S., Underhill, D. M., & Aderem, A. (2001). The innate immune response to bacterial flagellin is mediated by Toll-like receptor 5. Nature, 410(6832), 1099–1103. 10.1038/35074106

Hoffmann, S., Peterbauer, A., Schindler, S., Fennrich, S., Poole, S., Mistry, Y., Montag-Lessing, T., Spreitzer, I., Löschner, B., van Aalderen, M., Bos, R., Gommer, M., Nibbeling, R., Werner-Felmayer, G., Loitzl, P., Jungi, T., Brcic, M., Brügger, P., Frey, E., … Hartung, T. (2005). International validation of novel pyrogen tests based on human monocytoid cells. Journal of Immunological Methods, 298(1–2), 161–173. 10.1016/j.jim.2005.01.010

Huang, L.-Y., DuMontelle, J. L., Zolodz, M., Deora, A., Mozier, N. M., & Golding, B. (2009). Use of Toll-Like Receptor Assays To Detect and Identify Microbial Contaminants in Biological Products. Journal of Clinical Microbiology, 47(11), 3427–3434. 10.1128/JCM.00373-09

Kivioja, T., Vähärautio, A., Karlsson, K., Bonke, M., Enge, M., Linnarsson, S., & Taipale, J. (2012). Counting absolute numbers of molecules using unique molecular identifiers. Nature Methods, 9(1), 72–74. 10.1038/nmeth.1778

Klopfenstein, D. V., Zhang, L., Pedersen, B. S., Ramírez, F., Vesztrocy, A. W., Naldi, A., Mungall, C. J., Yunes, J. M., Botvinnik, O., Weigel, M., Dampier, W., Dessimoz, C., Flick, P., & Tang, H. (2018). GOATOOLS: A Python library for Gene Ontology analyses. Scientific Reports, 8(1). 10.1038/s41598-018-28948-z

Love, M. I., Huber, W., & Anders, S. (2014). Moderated estimation of fold change and dispersion for RNA-seq data with DESeq2. Genome Biology, 15(12), 550. 10.1186/s13059-014-0550-8

Molenaar-De Backer, M. W. A., Gitz, E., Dieker, M., Doodeman, P., & ten Brinke, A. (2021). Performance of monocyte activation test supplemented with human serum compared to fetal bovine serum. Altex, 38(2), 307–315. 10.14573/altex.2008261

Mueller, K., Chinchilla, D., Albert, M., Jehle, A. K., Kalbacher, H., Boller, T., & Felix, G. (2012). Contamination Risks in Work with Synthetic Peptides: flg22 as an Example of a Pirate in Commercial Peptide Preparations. The Plant Cell, 24(8), 3193–3197. 10.1105/tpc.111.093815

Muzellec, B., Teleńczuk, M., Cabeli, V., & Andreux, M. (2023). PyDESeq2: a python package for bulk RNA-seq differential expression analysis. Bioinformatics, 39(9), btad547. 10.1093/bioinformatics/btad547

Parra-Izquierdo, I., Lakshmanan, H. H. S., Melrose, A. R., Pang, J., Zheng, T. J., Jordan, K. R., Reitsma, S. E., McCarty, O. J. T., & Aslan, J. E. (2021). The Toll-Like Receptor 2 Ligand Pam2CSK4 Activates Platelet Nuclear Factor-κB and Bruton’s Tyrosine Kinase Signaling to Promote Platelet-Endothelial Cell Interactions. Frontiers in Immunology, 12. 10.3389/fimmu.2021.729951

Poltorak, A., He, X., Smirnova, I., Liu, M.-Y., Huffel, C. Van, Du, X., Birdwell, D., Alejos, E., Silva, M., Galanos, C., Freudenberg, M., Ricciardi-Castagnoli, P., Layton, B., & Beutler, B. (1998). Defective LPS Signaling in C3H/HeJ and C57BL/10ScCr Mice: Mutations in Tlr4 Gene. Science, 282(5396), 2085–2088. 10.1126/science.282.5396.2085

Salyer, A. C. D., & David, S. A. (2018). Transcriptomal signatures of vaccine adjuvants and accessory immunostimulation of sentinel cells by toll-like receptor 2/6 agonists. Human Vaccines and Immunotherapeutics, 14(7), 1686–1696. 10.1080/21645515.2018.1480284

Schwandner, R., Dziarski, R., Wesche, H., Rothe, M., & Kirschning, C. J. (1999). Peptidoglycan- and Lipoteichoic Acid-induced Cell Activation Is Mediated by Toll-like Receptor 2. Journal of Biological Chemistry, 274(25), 17406–17409. 10.1074/jbc.274.25.17406

Solati, S., Aarden, L., Zeerleder, S., & Wouters, D. (2015). An improved monocyte activation test using cryopreserved pooled human mononuclear cells. Innate Immunity, 21(7), 677–684. 10.1177/1753425915583365

Vapnik, V. N. (2000). The Nature of Statistical Learning Theory. Springer New York. 10.1007/978-1-4757-3264-1

Wolf, F. A., Angerer, P., & Theis, F. J. (2018). SCANPY: large-scale single-cell gene expression data analysis. Genome Biology, 19(1), 15. 10.1186/s13059-017-1382-0

