## Supplementary_Figures for "Identifying pyrogenic contaminants using transcriptomic profiling of monocyte activation test with machine learning"

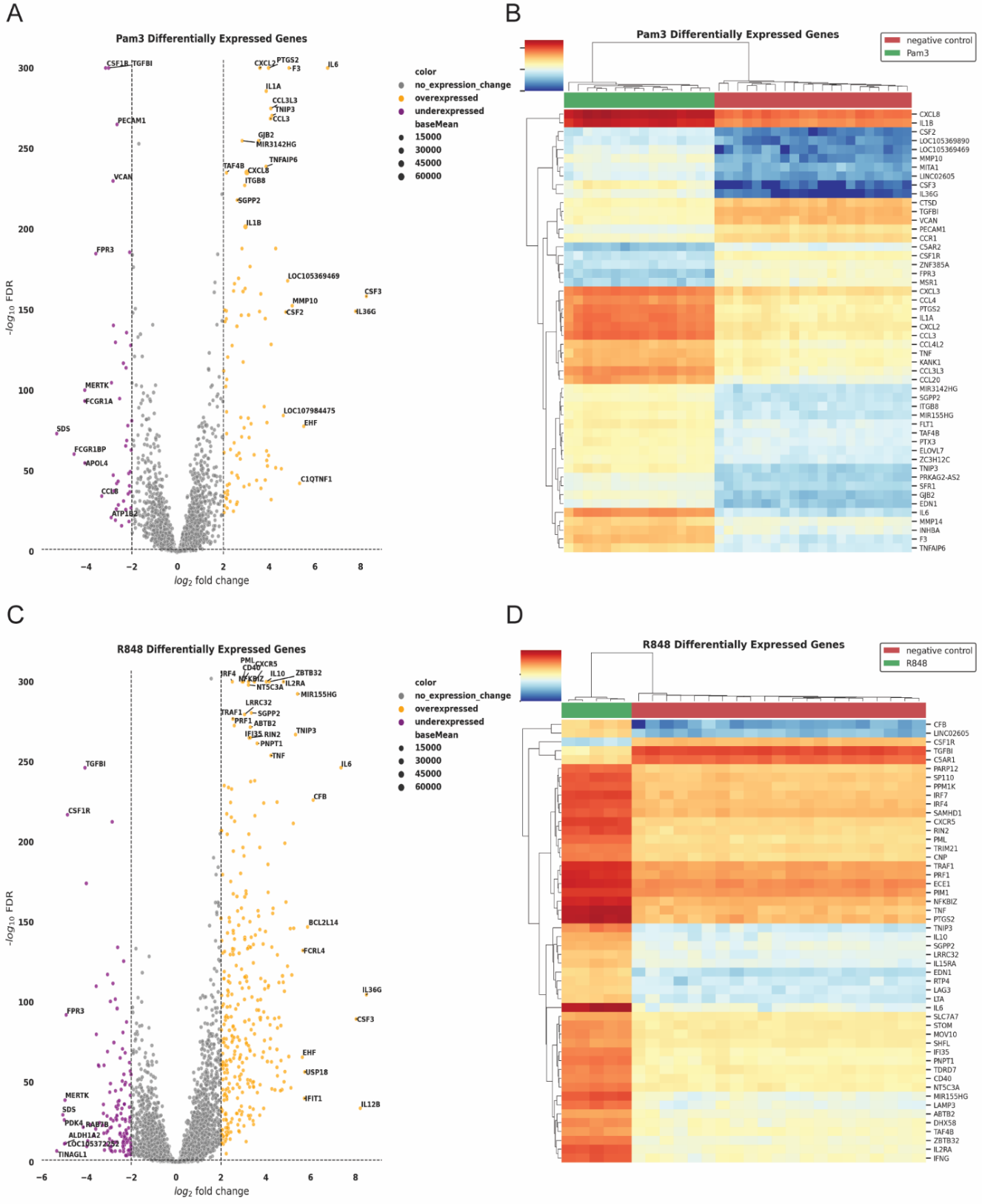

Supplementary Figure 1. Differential gene expression analysis for ligand-stimulated PBMCs. A) Up-and down-regulated genes for Pam3-stimulated PBMCs (FDR = 0.05, log2FoldChange =2), B) Differential gene expression for Pam3-stimulated monocytes. C) Up-and down-regulated genes for R848-stimulated PBMCs (FDR = 0.05, log2FoldChange =2), D) Differential gene expression for R848-stimulated PBMCs.

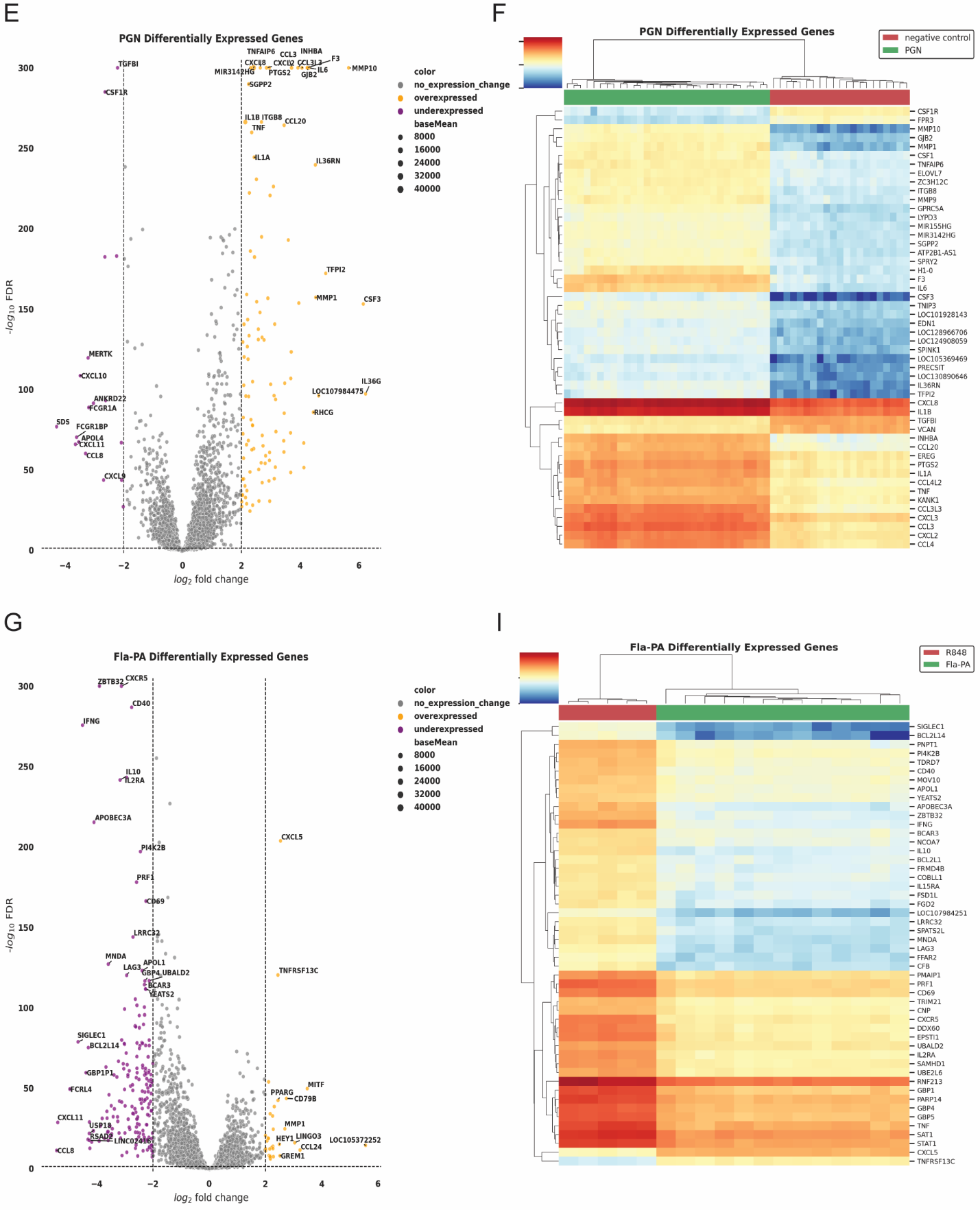

Supplementary Figure 1 continued. Differential gene expression analysis is for synthetic ligand-stimulated monocytes. E) Volcano plot shows up-and down-regulated genes for PGN-stimulated monocytes (FDR = 0.05, log2FoldChane =2), F) heatmap showing differential gene expression for PGN-stimulated monocytes. G) Volcano plot shows up-and down-regulated genes for Fla-PA-stimulated monocytes (FDR = 0.05, log2FoldChane =2), I) heatmap showing differential gene expression for Fla-PA-stimulated monocytes.

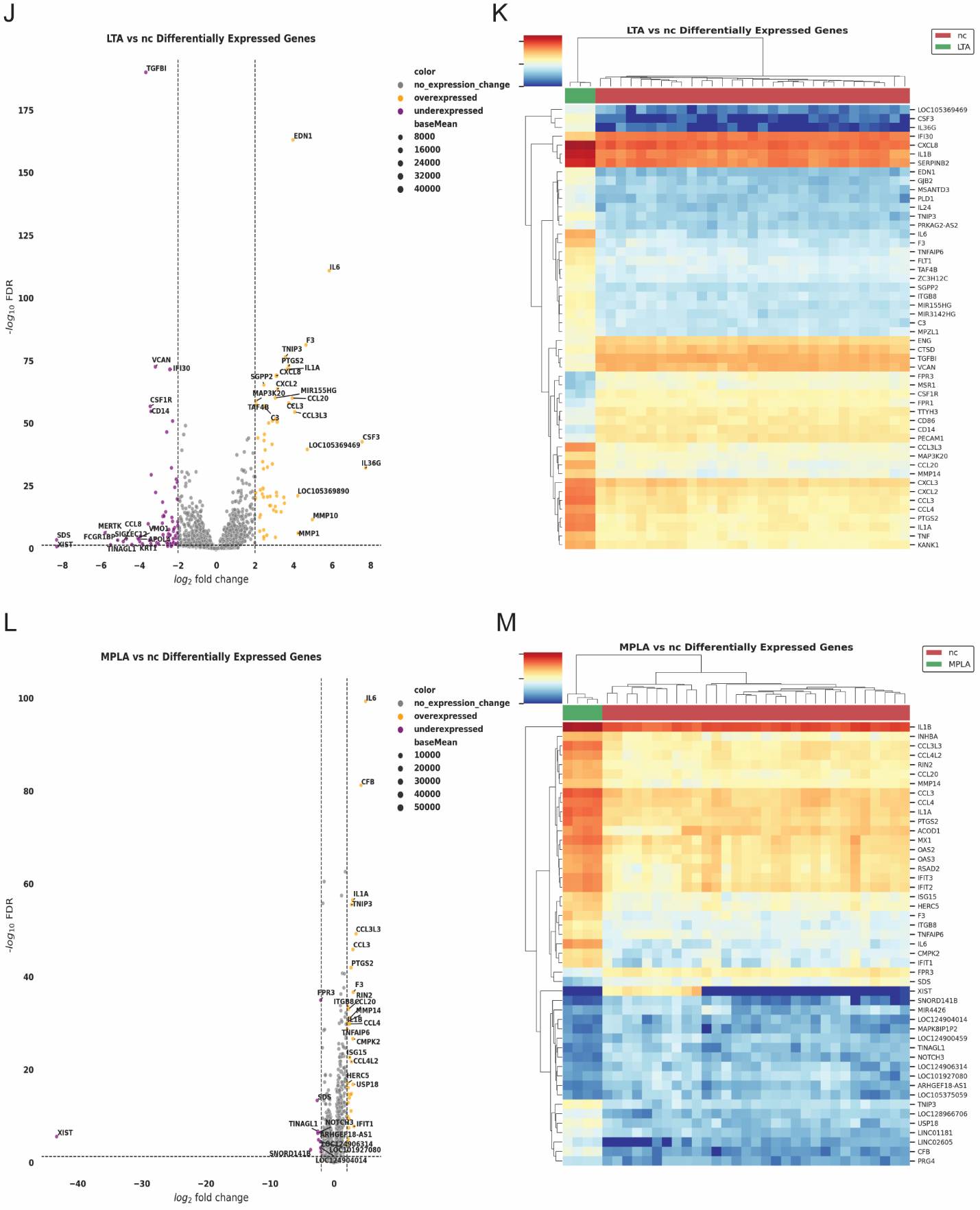

Supplementary Figure 1 continued. Differential gene expression analysis is for synthetic ligand-stimulated monocytes. J) Volcano plot shows up-and down-regulated genes for LTA-stimulated monocytes (FDR = 0.05, log2FoldChane =2), K) heatmap showing differential gene expression for LTA -stimulated monocytes. L) Volcano plot shows up-and down-regulated genes for MPLA-stimulated monocytes (FDR = 0.05, log2FoldChane =2), M) heatmap showing differential gene expression for MPLA -stimulated monocytes.

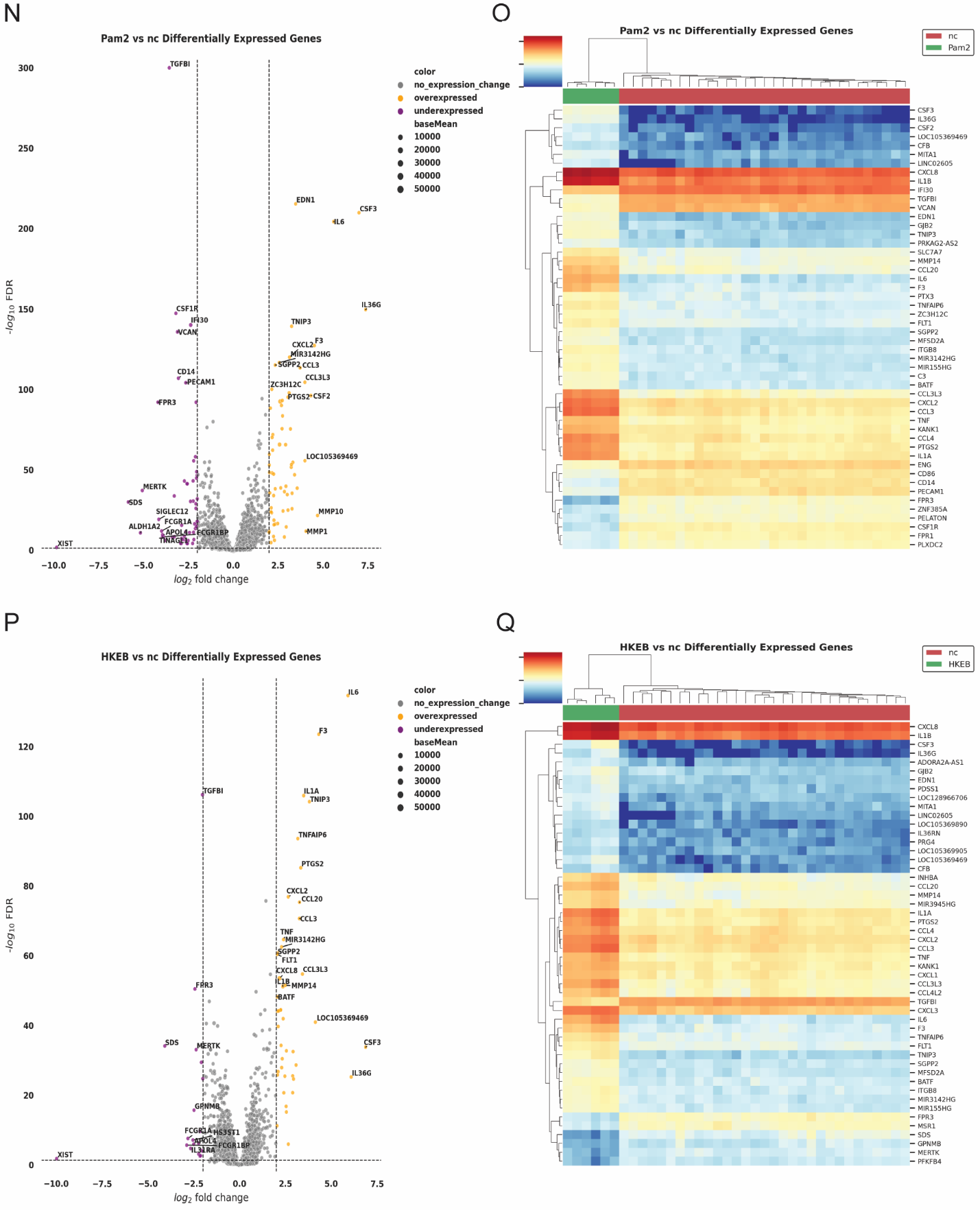

Supplementary Figure 1 continued. Differential gene expression analysis is for synthetic ligand-stimulated monocytes. N) Volcano plot shows up-and down-regulated genes for Pam2-stimulated monocytes (FDR = 0.05, log2FoldChane =2), O) heatmap showing differential gene expression for Pam2-stimulated monocytes. P) Volcano plot shows up-and down-regulated genes for heat-killed *E.coli-*stimulated monocytes (FDR = 0.05, log2FoldChane =2), Q) heatmap showing differential gene expression for heat-killed *E.coli* -stimulated monocytes.

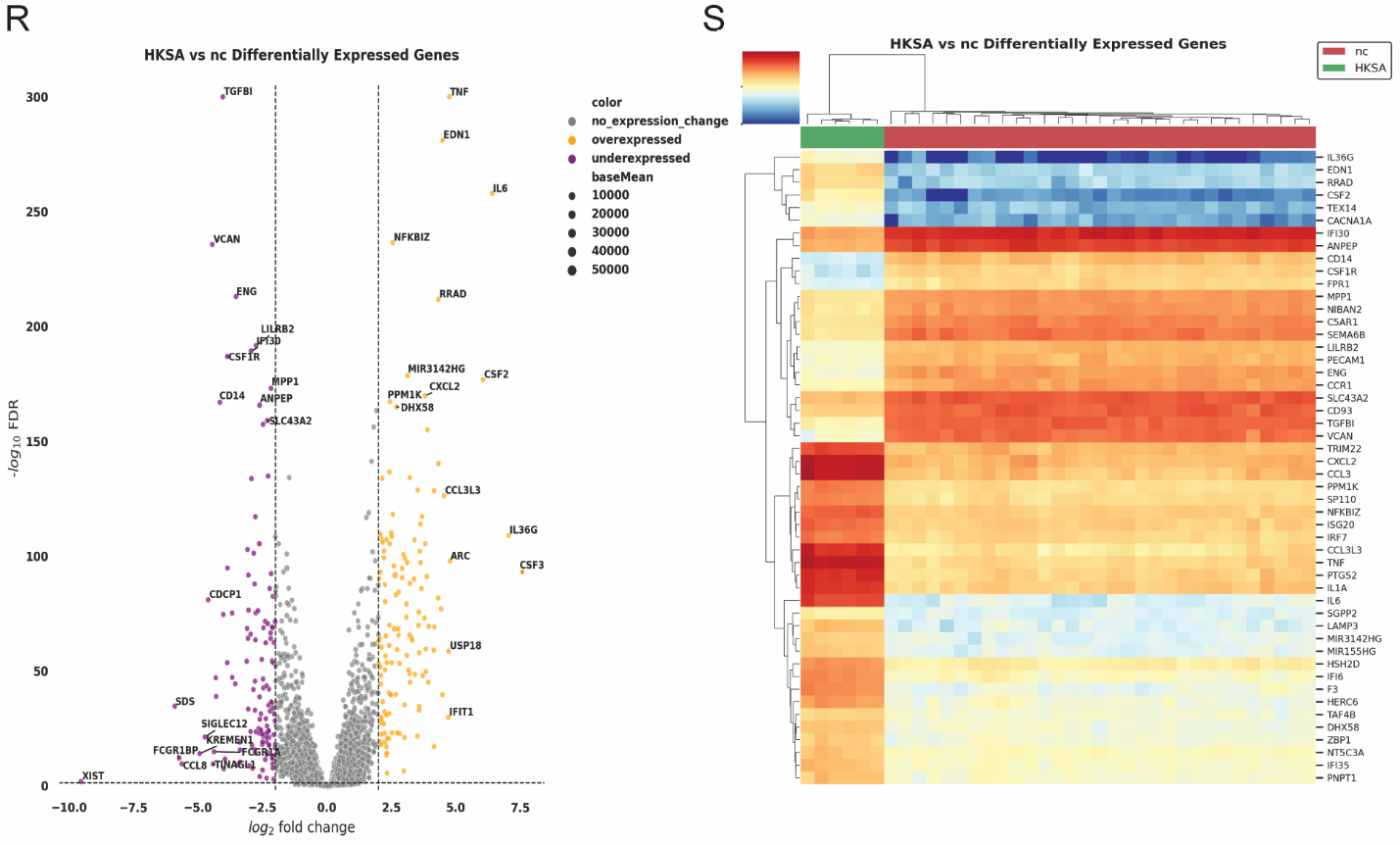
Supplementary Figure 1 continued. Differential gene expression analysis is for synthetic ligand-stimulated monocytes. R) Volcano plot shows up-and down-regulated genes for heat-killed *S. aureus-*stimulated monocytes (FDR = 0.05, log2FoldChane=2), S) heatmap showing differential gene expression for heat-killed *S. aureus*-stimulated monocytes.

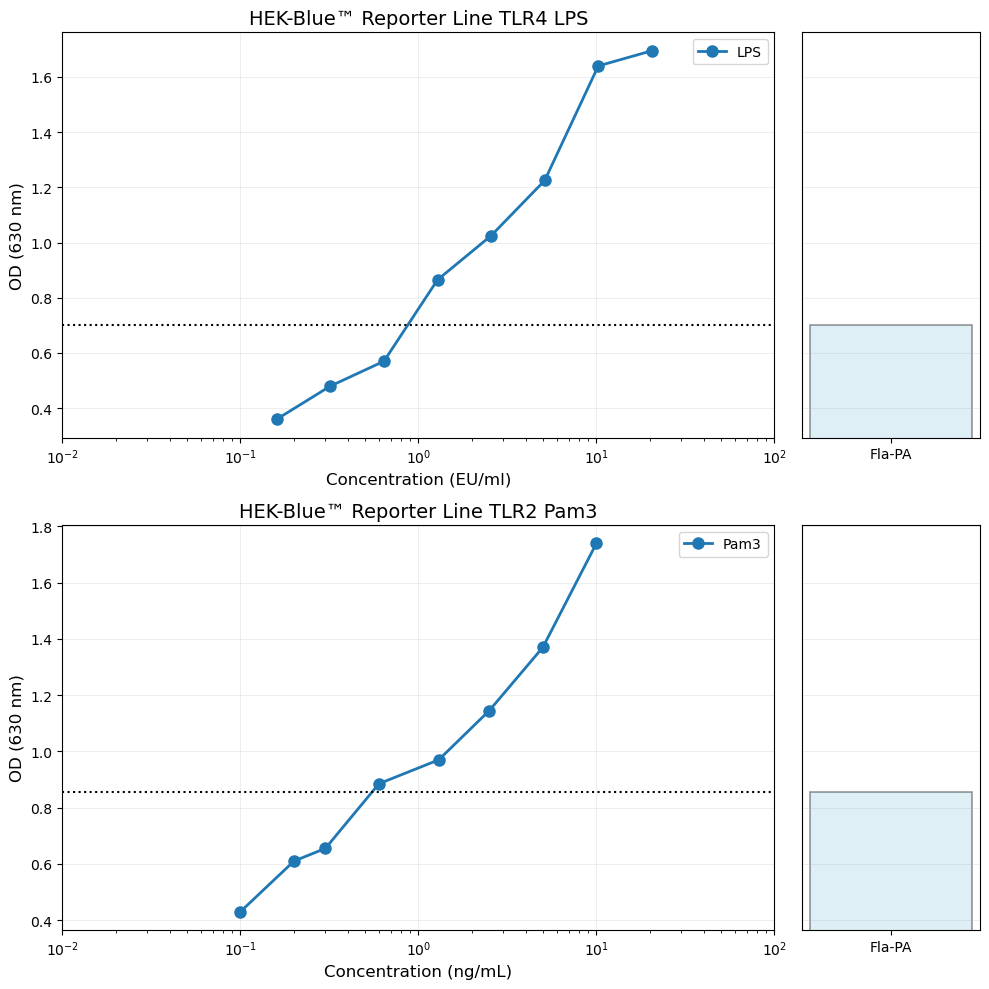

Supplementary Figure 2. HEK-Blue™ reporter cell line responses to LPS, Pam3, and Fla-PA. Top panel: HEK-Blue™ TLR4 response to LPS across a concentration range, with Fla-PA shown separately for comparison using the same y-axis. Bottom panel: HEK-Blue™ TLR2 response to Pam3 across a concentration range, with Fla-PA shown separately. Fla-PA was tested at 20 ng/ml, corresponding to the MAT signal equivalent of 0.16 EU/ml LPS. Ligand Pam3, PGN, R848 tested at 0.16 EU/ml equivalents were negative (below the standard curve) on both HEK-TLR4 and HEK-TLR2 cell lines.

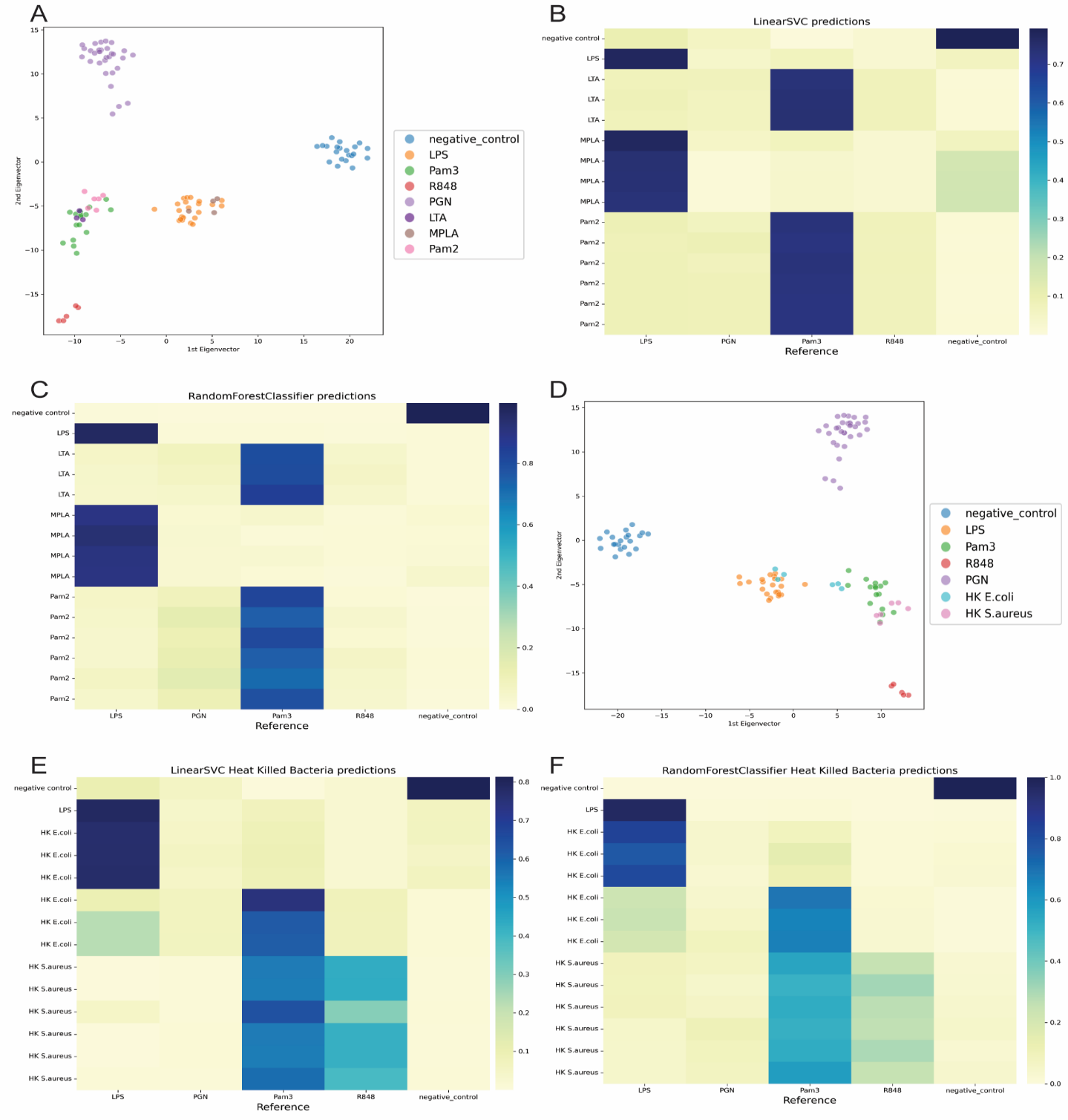
Supplementary Figure 3. ML model predictions on unseen TLR ligands and heat-killed bacterial samples after excluding Fla-PA from training. A) PCA of unseen ligands after feature selection, B,C) LinearSVC and RandomForest classification of unseen ligands, D) PCA of heat-killed E. coli and S. aureus after feature selection, E,F) LinearSVC and RandomForest classification of heat-killed samples.
